# Propagation mode shapes contrasting growth strategies through aquaporin networks in onion

**DOI:** 10.64898/2026.05.29.728701

**Authors:** Gloria Bárzana, Micaela Carvajal

## Abstract

Propagation mode strongly influences crop establishment, yet its impact on whole-plant transport strategies remains poorly understood. Here, we examined whether seed- and set-derived plants deploy contrasting aquaporin networks associated with different physiological behaviours in onion (*Allium cepa* L.). We combined genome-wide gene-family characterisation with transcriptomic, biochemical, and physiological analyses. Forty-eight aquaporin genes were identified and classified into four subfamilies (15 PIPs, 19 TIPs, 8 NIPs, and 6 SIPs), with evidence of lineage-specific expansion in the PIP1 and TIP2. Expression analyses revealed clear propagation-dependent patterns. Set-derived plants displayed higher expression of AcPIP1.1, several PIP2 isoforms and most TIP2 members in both roots and leaves, consistent with enhanced water and CO_2_ transport, higher stomatal conductance, transpiration, and photosynthetic rates. In contrast, seed-derived plants showed increased expression of specific PIPs, and several NIPs (AcNIP1.1, AcNIP3.1, AcNIP5.1, AcNIP5.2, and AcNIP2.1) associated with solute and H_2_O_2_ transport, coinciding with higher boron and hydrogen peroxide levels. These findings indicate that propagation origin is associated with alternative aquaporin-mediated transport strategies: set-derived plants favour a high-flux strategy that supports rapid growth, whereas seed-derived plants prioritize tighter internal regulation through solute redistribution and redox homeostasis. Our results provide a molecular-physiological framework linking propagation origin with resource-use strategies in onion.

**Highlight:** Propagation origin reprograms aquaporin expression in onion, generating contrasting hydraulic, metabolic and growth strategies.

**Graphical Abstract:** 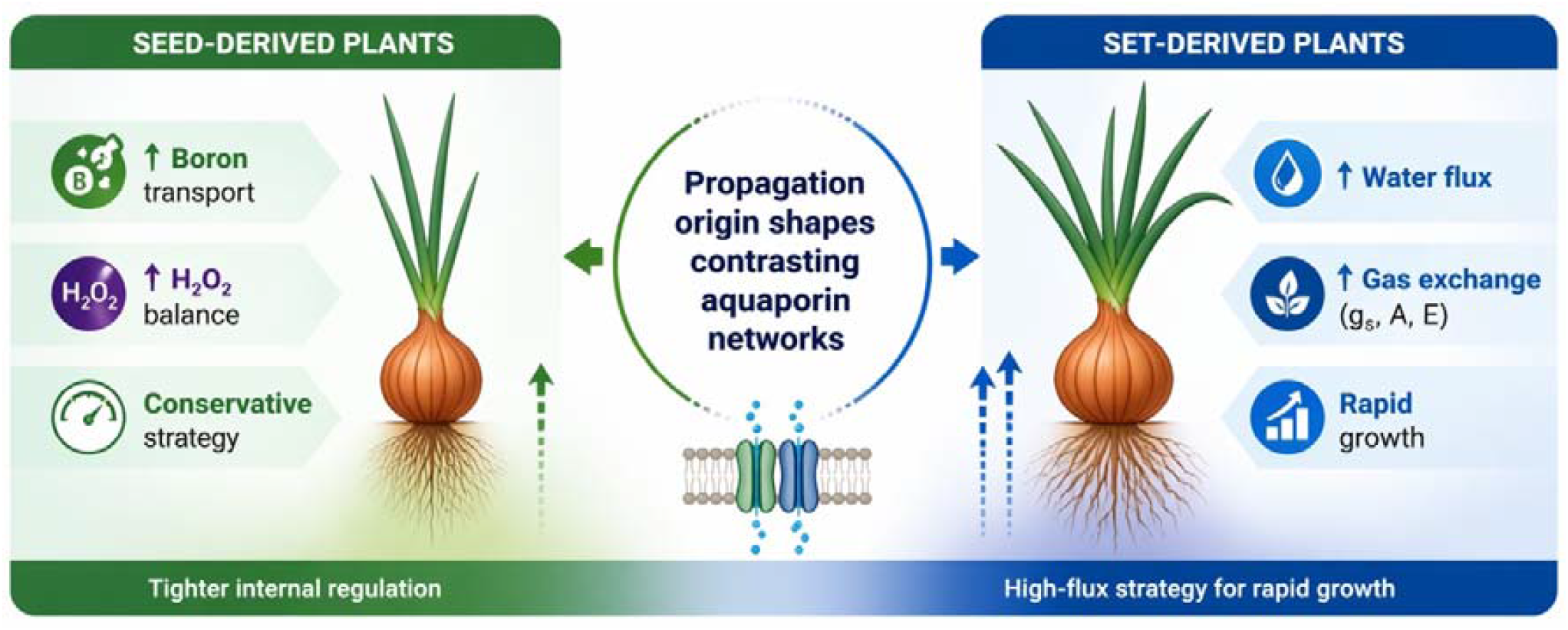

## 1. Introduction

Onion (*Allium cepa* L.) is one of the most widely cultivated vegetables globally (FAO, 2023). Belonging to the family Amaryllidaceae, onion is used in Mediterranean, Asian and African diets, valued for its well-known nutritional, culinary and medicinal properties (Brewster, 2008; Kuete, 2017; Kumar *et al*., 2022). The cultivation of onion is usually based on two propagation methods: seeds and small bulbs commonly known as onion sets. Seeds propagation is generally preferred because of its cost-effectiveness. They allow a higher planting density and therefore greater yield per unit area (Vojnović *et al*., 2023). However, onion seeds germinate slowly and are sensitive to environmental stress. In contrast, onion sets are pre-grown from seed, dried, and maintained in a dormant bulb stage, enabling rapid regrowth after planting (Pöhnl *et al*., 2019). Although using sets can increase production costs, farmers often use onion sets to minimize the impact of environmental stresses during the early stages of cultivation (Pöhnl *et al*., 2018). However, the contrasting developmental strategies on water and nutrient regulatory mechanisms that drive growth initiation in seed-propagated versus set-derived plants have not been studied.

Aquaporins (AQPs) are membrane intrinsic proteins (MIPs) that transport water and small solutes across biological membranes, playing central roles in osmoregulation, growth, reproduction and stress tolerance (Chaumont and Tyerman, 2017). In plants, AQPs are classified into five main subfamilies based on sequence homology and subcellular localisation: plasma membrane intrinsic proteins (PIPs), tonoplast intrinsic proteins (TIPs), nodulin-26-like intrinsic proteins (NIPs), small basic intrinsic proteins (SIPs), and the ancient X-intrinsic proteins (XIPs), which are absent in monocots such as onion (Bienert *et al*., 2011). These subfamilies differ in substrate specificity —from water and ammonia to metalloids or urea— and often show tissue-specific expression patterns, hinting at specialized physiological functions. The number of AQP isoforms varies across species —for example, 31 in melon (Lopez-Zaplana *et al*., 2020), 35 in arabidopsis (Johanson *et al*., 2001), 40 in rice (Raza *et al*., 2023), or 47 in tomato plants (Reuscher *et al*., 2013)— reflecting their species-specific adaptations and physiological needs. The tissue-specific expression of distinct AQP isoforms should be relevant in onion, where rapid growth from sets versus delayed establishment from seeds may rely on differential regulation of water and solute transport at the root and leaf levels. The study of aquaporins-mediated transport processes could provide molecular markers and targets for improving both propagation methods but, despite their relevance, no genome-wide analysis of AQP genes has yet been performed in this crop.

In that sense, although some initial steps have been taken in the molecular characterisation of onion [for review see (Sharma *et al*., 2024)], our understanding of the underlying genetic mechanisms remains limited. This gap may be due to the fact that the first high-quality onion genome assembly was not released until 2021 (Finkers *et al*., 2021). Based on a doubled-haploid line (DHCU066619) this genome has an estimated size of ~16⍰Gb, making it one of the largest among diploid cultivated species and particularly challenging due to its high repeat content and numerous pseudogenes (Finkers *et al*., 2021). In 2023, eight putative chromosomes were established, and their analysis suggests that the repetitive content may reflect adaptation to diverse environmental conditions (Hao *et al*., 2023). This genomic advance offers an unprecedented opportunity, although gene annotations remain inconsistent and require further refinement.

Understanding the molecular basis of water transport in onion involving aquaporins will provide crucial insights into its physiological plasticity driving different responses in seed-propagated versus set-derived plants. We hypothesize that the contrasting physiological performance observed between seed-derived and set-derived onion plants may be associated with distinct aquaporin expression patterns and subfamily-specific regulation. However, no systematic study has explored the composition, classification or expression dynamics of this gene family in *Allium*. To address this, we combined genomic, transcriptomic and physiological approaches to characterise aquaporin-mediated transport in *Allium cepa*. Specifically, we aim to (i) identify, analyse and characterise the AQP family in onion and (ii) link aquaporin expression patterns with propagation-dependent physiological performance.

## 2. Materials and methods

### 2.1 Plant material and experimental design

Twenty seeds of *Allium cepa* L. cv. A19 were sown in seedling trays containing TPS Dark Fine Low Fert substrate (Jiffy Products International B.V.). Trays were maintained in a controlled-environment chamber at 20–24 °C under a 14 h light/10 h dark photoperiod, 60–80% relative humidity, and 400 µmol m^−2^ s^−1^ photosynthetically active radiation (PAR). During early establishment, plants were irrigated with Hoagland nutrient solution to ensure adequate water and nutrient supply. Forty-five days after sowing, seed-derived seedlings were trimmed to standardize shoot length and reduce developmental variability among individuals without affecting subsequent regrowth. Thereafter, irrigation was switched to water only in order to match the management conditions used for onion sets. Onion sets of the same cultivar were planted in the same substrate and maintained in parallel under identical chamber conditions. To compare propagation systems at equivalent developmental stage rather than chronological age, sets were planted later so that both seed-derived and set-derived plants reached similar shoot height and the 3–4 true-leaf stage simultaneously. Plants were then sampled in parallel for all subsequent analyses.

### 2.2 Physiological measurements

Physiological measurements were performed between 09:00 and 11:00 immediately before harvest on fully expanded young leaves from 10 biological replicates per treatment. Gas-exchange parameters were measured using a CIRAS-3 portable infrared gas analyser (PP Systems, Amesbury, MA, USA). Chamber conditions were set to 400 µmol mol^−1^ CO_2_, temperature 22–25 °C, relative humidity 40–60%, and irradiance of 600 µmol photons m^−2^ s^−1^. Net photosynthetic rate (A), stomatal conductance (gs), transpiration rate (E), incident photosynthetically active radiation (PAR), and intercellular CO_2_ concentration (Ci) were recorded directly by the instrument. Photosynthetic CO_2_ assimilation (A/PAR) and instantaneous water use efficiency (WUE, A/E) were calculated according to Hu et al. (2010) and apparent carboxylation efficiency (ACE, A/Ci) according to Hornyák et al. (2005). Fresh weight (FW) of shoots and roots was determined immediately after harvest.

### 2.4 Biochemical analyses

Hydrogen peroxide (H_2_O_2_) content was determined in six biological replicates per treatment following the method of Velikova et al. (2000). Absorbance was measured using a PowerWave XS2 spectrophotometer (BioTek Instruments, Winooski, VT, USA), and H_2_O_2_ concentration was calculated from the corresponding extinction coefficient as described in the original method. For boron determination, finely ground dry tissue samples were digested in HNO_3_:HClO_4_ (2:1, v/v). Boron concentration was quantified in triplicate using an ICP emission spectrophotometer (Iris Intrepid II, Thermo Electron Corporation, Franklin, TN, USA). Results were expressed as mg kg^−1^ dry weight. Individual biological replicate values are provided in Supplementary Table S1.

### 2.5 Identification of putative Allium cepa aquaporins (AcAQPs)

To comprehensively identify aquaporin genes in *Allium cepa*, we combined homology-based searches and *ab initio* prediction methods. The complete set of aquaporin sequences of *Allium cepa* was compiled from multiple sources: 79 non-curated sequences were obtained from aquaporins annotated in AlliumDB database (http://bioinformatics.nat300.top/Allium/) (Yang *et al*., 2024), 49 non-curated sequences from aquaporins annotated in NABIC database (https://nabic.rda.go.kr), and 117 non-curated sequences were obtained by tBLASTn searches using all *Arabidopsis thaliana* L. (Johanson *et al*., 2001) and *Cucumis melo* L. (Lopez-Zaplana *et al*., 2020) aquaporins against whole *Allium cepa* genome sequences (ASM3076508v1 and DHCU066619 genomes) obtained from NCBI using Geneious Prime software. All sequences were individually analysed. Gene models were predicted using MAKER, GFFreader (via Galaxy platform) and FGENESH (SoftBerry) to reconstruct the coding sequences. Each predicted protein was validated by BLASTp against the NCBI nr protein database and aligned using Clustal Omega to detect redundancies and structural completeness. Based on these analyses, sequences were classified as full-length aquaporins, partial sequences or pseudogenes, and aquaporin-related genomic fragments. Finally, only full-length sequences and partial sequences with transcriptional support were selected for subsequent analysis.

### 2.5 Phylogenetic Classification and Nomenclature

Curated full-length aquaporin proteins were named according to phylogenetic relationships with reference aquaporins from *Arabidopsis thaliana* L., *Zea mays* L. and *Oryza sativa* L. (Johanson *et al*., 2001; Su *et al*., 2022; Raza *et al*., 2023). Phylogenetic analysis was performed using MEGA12 software (version 12). Multiple sequence alignment of the deduced amino acid sequences was conducted using MUSCLE with default parameters. Phylogenetic reconstruction was conducted using the Maximum Likelihood method with the JTT substitution model and a discrete Gamma distribution (five categories; Γ = 1.0). Node support was assessed by 1000 bootstrap replicates. Phylogenetic trees were visualized using iTOL.

### 2.6 Gene sequences analysis

Conserved motif identification was performed using MEME Suite v5.5.8. Exon–intron organization was analysed using GSDS v2.0 (Hu et al. 2015a). Chromosomal localisation of aquaporin genes was retrieved from AlliumDB.

### 2.7 Protein characterisation

Protein features such as amino acid number, isoelectric point (pI), molecular weight (Mw), instability index, aliphatic index, and GRAVY score, were calculated using ProtParam (ExPASy). Transmembrane helices were predicted using Protter (https://wlab.ethz.ch/protter/start/) and TMHMM v2.0. (https://services.healthtech.dtu.dk/) Subcellular localisation was predicted using DeepLoc and cross-validated with Plant-mPLoc (http://www.csbio.sjtu.edu.cn/bioinf/plant-multi/). Data are presented in Table 1 and Supplementary Table S2. Three-dimensional structures were predicted using Phyre2 (https://www.sbg.bio.ic.ac.uk/phyre2) for aquaporins with uncertain structural completeness.

**Table 1.**
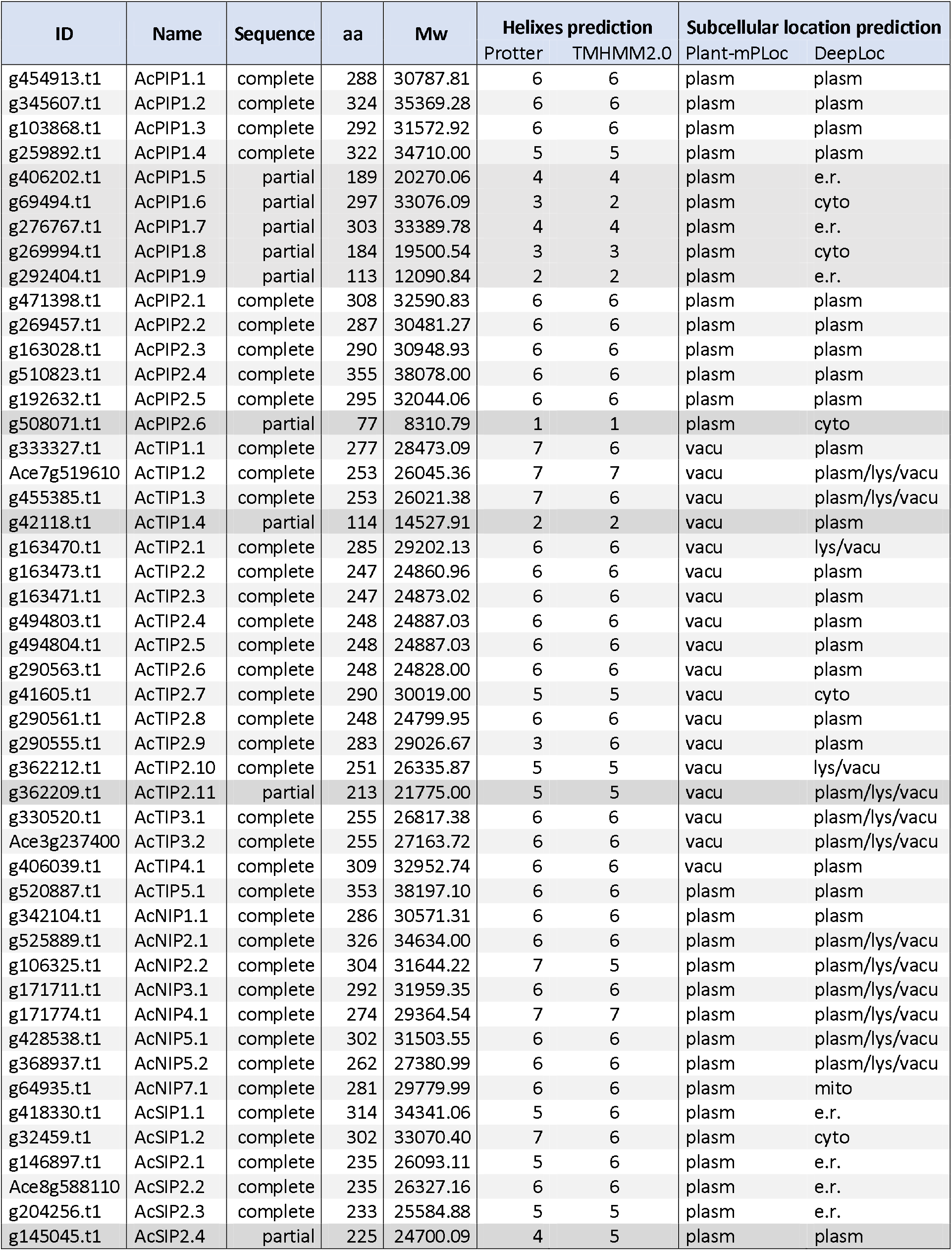
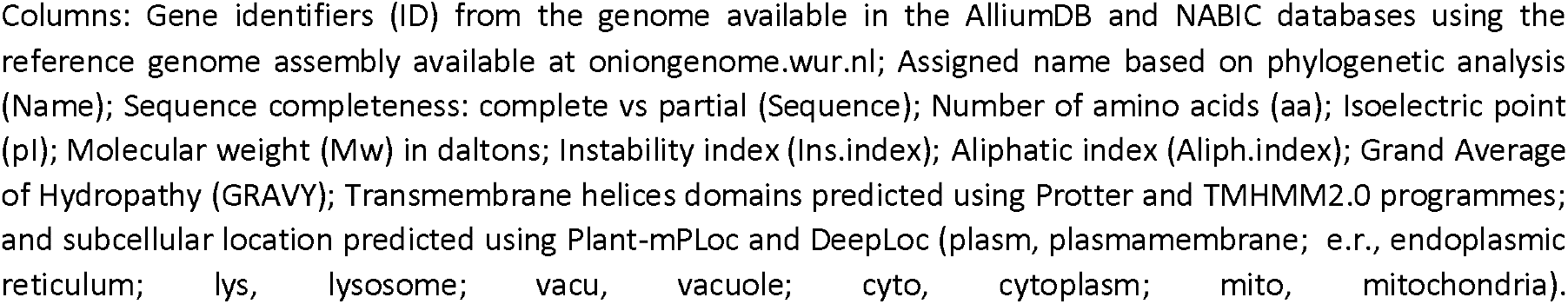
Structural and physicochemical characteristics of aquaporins identified in *Allium cepa* L. (AcAQPs).

| ID | Name | Sequence | aa | Mw | Helixes prediction |  | Subcellular location prediction |  |
| --- | --- | --- | --- | --- | --- | --- | --- | --- |
|  |  |  |  |  | Protter | TMHMM2.0 | Plant-mPloc | DeepLoc |
| g454913.t1 | AcPIP1.1 | complete | 288 | 30787.81 | 6 | 6 | plasm | plasm |
| g345607.t1 | AcPIP1.2 | complete | 324 | 35369.28 | 6 | 6 | plasm | plasm |
| g103868.t1 | AcPIP1.3 | complete | 292 | 31572.92 | 6 | 6 | plasm | plasm |
| g259892.t1 | AcPIP1.4 | complete | 322 | 34710.00 | 5 | 5 | plasm | plasm |
| g406202.t1 | AcPIP1.5 | partial | 189 | 20270.06 | 4 | 4 | plasm | e.r. |
| g69494.t1 | AcPIP1.6 | partial | 297 | 33076.09 | 3 | 2 | plasm | cyto |
| g276767.t1 | AcPIP1.7 | partial | 303 | 33389.78 | 4 | 4 | plasm | e.r. |
| g269994.t1 | AcPIP1.8 | partial | 184 | 19500.54 | 3 | 3 | plasm | cyto |
| g292404.t1 | AcPIP1.9 | partial | 113 | 12090.84 | 2 | 2 | plasm | e.r. |
| g471398.t1 | AcPIP2.1 | complete | 308 | 32590.83 | 6 | 6 | plasm | plasm |
| g269457.t1 | AcPIP2.2 | complete | 287 | 30481.27 | 6 | 6 | plasm | plasm |
| g163028.t1 | AcPIP2.3 | complete | 290 | 30948.93 | 6 | 6 | plasm | plasm |
| g510823.t1 | AcPIP2.4 | complete | 355 | 38078.00 | 6 | 6 | plasm | plasm |
| g192632.t1 | AcPIP2.5 | complete | 295 | 32044.06 | 6 | 6 | plasm | plasm |
| g508071.t1 | AcPIP2.6 | partial | 77 | 8310.79 | 1 | 1 | plasm | cyto |
| g333327.t1 | AcTIP1.1 | complete | 277 | 28473.09 | 7 | 6 | vacu | plasm |
| Ace7g519610 | AcTIP1.2 | complete | 253 | 26045.36 | 7 | 7 | vacu | plasm/lys/vacu |
| g455385.t1 | AcTIP1.3 | complete | 253 | 26021.38 | 7 | 6 | vacu | plasm/lys/vacu |
| g42118.t1 | AcTIP1.4 | partial | 114 | 14527.91 | 2 | 2 | vacu | plasm |
| g163470.t1 | AcTIP2.1 | complete | 285 | 29202.13 | 6 | 6 | vacu | lys/vacu |
| g163473.t1 | AcTIP2.2 | complete | 247 | 24860.96 | 6 | 6 | vacu | plasm |
| g163471.t1 | AcTIP2.3 | complete | 247 | 24873.02 | 6 | 6 | vacu | plasm |
| g494803.t1 | AcTIP2.4 | complete | 248 | 24887.03 | 6 | 6 | vacu | plasm |
| g494804.t1 | AcTIP2.5 | complete | 248 | 24887.03 | 6 | 6 | vacu | plasm |
| g290563.t1 | AcTIP2.6 | complete | 248 | 24828.00 | 6 | 6 | vacu | plasm |
| g41605.t1 | AcTIP2.7 | complete | 290 | 30019.00 | 5 | 5 | vacu | cyto |
| g290561.t1 | AcTIP2.8 | complete | 248 | 24799.95 | 6 | 6 | vacu | plasm |
| g290555.t1 | AcTIP2.9 | complete | 283 | 29026.67 | 3 | 6 | vacu | plasm |
| g362212.t1 | AcTIP2.10 | complete | 251 | 26335.87 | 5 | 5 | vacu | lys/vacu |
| g362209.t1 | AcTIP2.11 | partial | 213 | 21775.00 | 5 | 5 | vacu | plasm/lys/vacu |
| g330520.t1 | AcTIP3.1 | complete | 255 | 26817.38 | 6 | 6 | vacu | plasm/lys/vacu |
| Ace3g237400 | AcTIP3.2 | complete | 255 | 27163.72 | 6 | 6 | vacu | plasm/lys/vacu |
| g406039.t1 | AcTIP4.1 | complete | 309 | 32952.74 | 6 | 6 | vacu | plasm |
| g520887.t1 | AcTIP5.1 | complete | 353 | 38197.10 | 6 | 6 | plasm | plasm |
| g342104.t1 | AcNIP1.1 | complete | 286 | 30571.31 | 6 | 6 | plasm | plasm |
| g525889.t1 | AcNIP2.1 | complete | 326 | 34634.00 | 6 | 6 | plasm | plasm/lys/vacu |
| g106325.t1 | AcNIP2.2 | complete | 304 | 31644.22 | 7 | 5 | plasm | plasm/lys/vacu |
| g171711.t1 | AcNIP3.1 | complete | 292 | 31959.35 | 6 | 6 | plasm | plasm/lys/vacu |
| g171774.t1 | AcNIP4.1 | complete | 274 | 29364.54 | 7 | 7 | plasm | plasm/lys/vacu |
| g428538.t1 | AcNIP5.1 | complete | 302 | 31503.55 | 6 | 6 | plasm | plasm/lys/vacu |
| g368937.t1 | AcNIP5.2 | complete | 262 | 27380.99 | 6 | 6 | plasm | plasm/lys/vacu |
| g64935.t1 | AcNIP7.1 | complete | 281 | 29779.99 | 6 | 6 | plasm | mito |
| g418330.t1 | AcSIP1.1 | complete | 314 | 34341.06 | 5 | 6 | plasm | e.r. |
| g32459.t1 | AcSIP1.2 | complete | 302 | 33070.40 | 7 | 6 | plasm | cyto |
| g146897.t1 | AcSIP2.1 | complete | 235 | 26093.11 | 5 | 6 | plasm | e.r. |
| Ace8g588110 | AcSIP2.2 | complete | 235 | 26327.16 | 6 | 6 | plasm | e.r. |
| g204256.t1 | AcSIP2.3 | complete | 233 | 25584.88 | 5 | 5 | plasm | e.r. |
| g145045.t1 | AcSIP2.4 | partial | 225 | 24700.09 | 4 | 5 | plasm | plasm |

Columns: Gene identifiers (ID) from the genome available in the AlliumDB and NABIC databases using the reference genome assembly available at oniongenome.wur.nl; Assigned name based on phylogenetic analysis (Name); Sequence completeness: complete vs partial (Sequence); Number of amino acids (aa); Isoelectric point (pI); Molecular weight (Mw) in daltons; Instability index (Ins.index); Aliphatic index (Aliph.index); Grand Average of Hydropathy (GRAVY); Transmembrane helices domains predicted using Protter and TMHMM2.0 programmes; and subcellular location predicted using Plant-mPLOC and DeepLoc (plasm, plasmamembrane; e.r., endoplasmic reticulum; lys, lysosome; vacu, vacuole; cyto, cytoplasm; mito, mitochondria).

### 2.8 Solutes transport prediction

Alignment with representative aquaporins transporting H_2_O (SoPIP2.1) (Törnroth-Horsefield *et al*., 2006), ions (OsPIP2.4) (Ono *et al*., 2025; Qiu *et al*., 2025), CO_2_ (AtPIP2.1) (Chen *et al*., 2023), Urea (AtTIP5.1) (Dynowski *et al*., 2008*a*), NH_3_ (AtTIP2.1) (Kirscht *et al*., 2016), Boron (AtNIP5.1), silicon (OsNIP2.1) (Mitani-Ueno *et al*., 2011; Van den Berg *et al*., 2021) and glycerol (LIMP2) (Wallace *et al*., 2002) were conducted using Clustal Omega v1.2.4 from EMBL’s European Bioinformatics Institute (EMBL-EBI) server. Conserved residues associated with substrate selectivity, including the NPA motifs, aromatic/arginine (Ar/R) filter, and Froger positions, were manually inspected from the alignments (Froger *et al*., 1998; Murata *et al*., 2000; Sui *et al*., 2001). Putative transport capacities were inferred by comparison with arabidopsis, rice and maize orthologues together with published mutational, structural, and functional evidence, following the framework described by López-Zaplana et al. 2020 (2020). For isoforms showing unusual residues in the Ar/R filter or Froger positions, 3D models were generated using SWISS-MODEL. Predicted pore architecture and structural features were further inspected using Mol*. These predictions should be considered hypothesis-generating and require experimental validation.

### 2.9 Public RNA-seq expression profiling

Publicly available onion RNA-seq datasets were retrieved from the NCBI Sequence Read Archive (NCBI-SRA) to assess tissue-specific AcAQP expression patterns. Only control samples from plants older than 3 months and with biological replication were included. Raw reads (SRR, DRR, and ERR accessions) were downloaded from the European Nucleotide Archive and processed in Galaxy Europe. BioProject accessions are listed in Supplementary Table S3. Read quality was assessed using Falco. Filtered reads were aligned to the *Allium cepa* reference genome (oniongenome.wur.nl) using HISAT2, and gene-level counts were generated with featureCounts. Expression matrices were normalized using DESeq2 with variance stabilizing transformation (VST). Principal component analysis of VST-transformed data was used to evaluate sample clustering and potential batch effects among BioProjects (Supplementary Fig. S1). Tissue-specific aquaporin expression patterns were visualized as z-score heatmaps using OriginPro 2023 (OriginLab, Northampton, MA, USA).

### 2.10 Experimental RNA-seq analysis

For comparative transcriptomic analysis, six biological replicates per treatment (seed-derived and set-derived seedlings) were harvested at the matched developmental stage. Leaves and roots were collected separately. Total RNA was extracted using the NZY Total RNA Isolation kit (Nzytech, Lisbon, Portugal) and RNA integrity was assessed by RIN analysis (Schroeder *et al*., 2006) using an Agilent 2100 Bioanalyzer (Agilent Technologies, Santa Clara, CA, USA) with the 2100 Expert Eukaryote Total RNA Nano chip. cDNA libraries were prepared using the Illumina Stranded mRNA Prep, Ligation kit (Illumina Inc., San Diego, CA, USA) and quantified using a Qubit 4 Fluorometer (Thermo Fisher Scientific, Waltham, MA, USA). Library quality was further validated using the Agilent 2100 Bioanalyzer with the High Sensitivity DNA kit (Agilent Technologies). Sequencing was performed on an Illumina NextSeq 1000/2000 platform (100 bp paired-end reads; minimum 25 million reads per sample) with P3 flow cell chemistry (200 cycles). Raw reads were trimmed using fastp v0.20.0 (Chen *et al*., 2018) and pseudoaligned to the *Allium cepa* reference transcriptome using Salmon v0.13.1 (Patro *et al*., 2017). Transcript abundances were summarized to gene level using tximport. Log2FoldChange values and Normalized expression values for aquaporin genes were obtained using the variance stabilizing transformation (VST) in DESeq2 (Love *et al*., 2014). Heatmaps were generated using OriginPro 2023b (OriginLab Corporation, Northampton, MA, USA).

### 2.11 Statistical analysis

Physiological and biochemical measurements were obtained from independent biological replicates, as indicated in each subsection. Gas-exchange and fresh-weight parameters were measured in ten biological replicates per propagation mode, whereas H_2_O_2_ content was determined in six biological replicates per propagation mode. Boron concentration was measured in triplicate analytical determinations from finely ground dry tissue samples. Data are presented as mean ± standard error (SE). Differences between seed-derived and set-derived plants were evaluated by one-way analysis of variance (ANOVA) followed by Duncan’s multiple range post hoc test (p < 0.05). The assumptions of normality and homogeneity of variance were checked before ANOVA. For RNA-seq analyses, six biological replicates per propagation mode and tissue were used. Differential gene expression was performed using DESeq2, applying its internal normalization procedure and Wald statistics. Variance stabilizing transformation (VST) was used for exploratory analyses, PCA and heatmap visualization. PCA and graphical representations were generated using OriginPro 2023/2023b (OriginLab, Northampton, MA, USA). Statistical analyses were performed using SPSS v29.0 (IBM Corp., Armonk, NY, USA), DESeq2 and OriginPro.

## 3. Results

### 3.1 Identification of aquaporin genes

By combining homology-based searches from the AlliumDB and NABIC databases with ab initio prediction methods, 243 candidate sequences were retrieved from Allium cepa. Of these, 57 showed similarity to aquaporins, including 39 full-length putative functional aquaporins and 18 partial or pseudogenic sequences, whereas the remaining candidates were discarded as aquaporin-related genomic fragments lacking key functional domains.

Several candidate aquaporins displayed atypical transmembrane predictions despite containing conserved aquaporin motifs and appropriate sequence lengths (Table 1). Structural modelling confirmed the canonical aquaporin fold and supported their classification as functional aquaporins (Supplementary Fig. S2). Motif analysis (Supplementary Fig. S3A) revealed that in g259892.t1 and g362212.t1 genes motif 6 was not detected while in g41605.t1 motif 7 was missing. Furthermore, transcriptional evidence supported their classification as fully functional aquaporins. The sequence g406202.t1 was classified as a partial aquaporin due to the absence of a central region in all gene model predictions despite showing transcriptional support.

Given the highly repetitive nature of the onion genome and the likelihood of assembly fragmentation (Finkers *et al*., 2021), partial sequences with transcriptional evidence were retained as “partial” aquaporins (Table 1). Altogether, 48 AcAQP isoforms were identified and included in subsequent analyses.

### 3.2 Phylogenetic analysis and classification

Phylogenetic analysis grouped the 48 AcAQPs into the four canonical plant aquaporin subfamilies (Fig. 1): fifteen PIPs, nineteen TIPs, eight NIPs and six SIPs. Gene nomenclature was assigned according to phylogenetic relationships with Arabidopsis, rice and maize aquaporins (Fig. 1). Names assigned to each isoform are presented in Table 1. Motif composition and exon–intron organization were highly conserved within subfamilies and consistent with patterns reported in other angiosperms (Supplementary Fig. S3-S4). The most conserved motifs across all aquaporins were motifs 1 and 5 corresponding to NPA motif and the transmembrane helix 3, two highly conserved structural components in evolution (Kruse *et al*., 2006; Israel *et al*., 2023). Based on current genome annotations, only 17 aquaporin genes could be assigned to chromosomes, whereas the remaining genes were located on unplaced scaffolds (Supplementary Table S4).

**Fig. 1.**
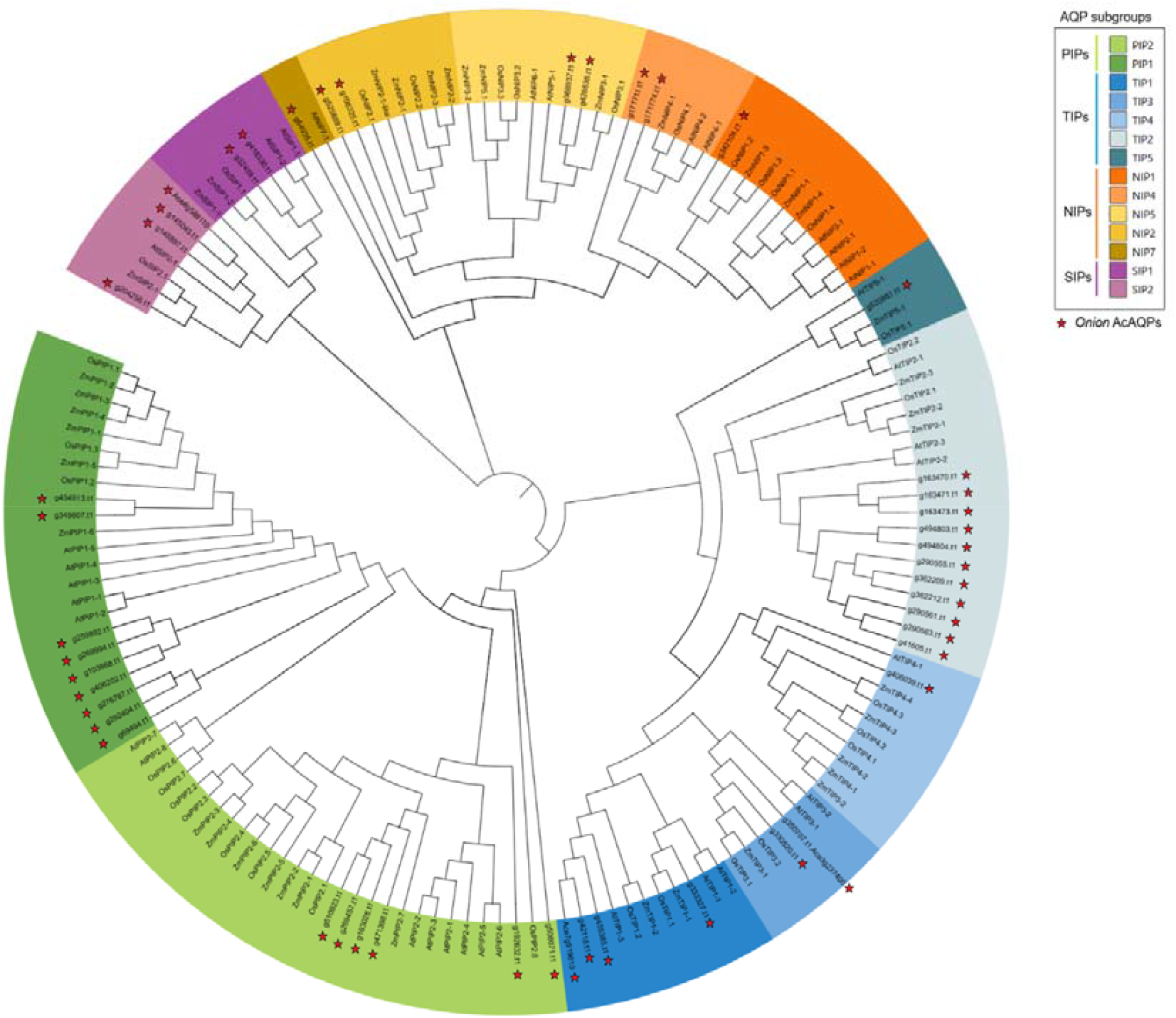
Phylogenetic relationships among onion aquaporins and reference aquaporins from Arabidopsis thaliana (AtAQP), Oryza sativa (OsAQP), and Zea mays (ZmAQP). Alignment was done with MUSCLE algorithm using the Maximum Likelihood method with the JTT matrix-based model and a discrete Gamma distribution to model evolutionary rate differences among sites (5 categories, Gamma = 1.0). The tree design was carried out using iTOL Interactive Tree of Life online tool. The different aquaporin subfamilies are highlighted in different colours: green for PIPs (green), TIPs (blue), NIPs (orange), and SIPs (purple). Different shades indicate subgroups within each subfamily. Red stars indicate Allium cepa aquaporins identified in this study. **Alt text:** Circular phylogenetic tree showing aquaporin protein sequences from onion together with reference aquaporins from Arabidopsis, rice, and maize. Branches are grouped into four colour-coded aquaporin subfamilies: PIPs (green), TIPs (blue), NIPs (orange), and SIPs (purple), with different shades indicating subgroups within each family. Onion aquaporins identified in this study are marked with red stars.

### 3.3 Protein sequences analysis and subcellular location prediction

All physical characteristics of the putative *Allium cepa* aquaporins are summarized in Table 1. The identified putative aquaporins comprised between 233 and 355 amino acids, which is the standard size for this protein family (Johanson *et al*., 2001). A few exceptions corresponded to shorter partial sequences described above as AcPIP2.6 with only 77 aa, AcPIP1.5 with 189 aa, AcPIP1.8 with 184 aa, AcPIP1.9 with 113 aa, AcTIP1.4 with 114, AcTIP2.11 with 213 and AcSIP2.4 with 225 (Table 1). Physicochemical properties were consistent with those reported for aquaporins from other plant species (Supplementary Table S2) (Johanson *et al*., 2001; Hu *et al*., 2015; Lopez-Zaplana *et al*., 2020). Molecular weight, instability index, aliphatic index and GRAVY values fell within the ranges previously reported for plant aquaporins (Hu *et al*., 2015; Qian *et al*., 2019; Lopez-Zaplana *et al*., 2020; Malik *et al*., 2023)

Regarding subcellular location, most isoforms were predicted in their expected locations (Maurel *et al*., 2008) with PIPs assigned to the plasma membrane, TIPs to the tonoplast, and SIPs predominantly to the endoplasmic reticulum. In the case of NIPs, they were mostly recognized in plasma membrane, only AcNIP7.1 was assigned to mitochondria by DeepLoc (Table 1).

### 3.4 Solute transport prediction

In order to locate the key residues related to the aquaporin selectivity presented in Table 2, we aligned the AcAQPs sequences with well-characterised transporters of H_2_O_2_ (SoPIP2.1) (Törnroth-Horsefield *et al*., 2006), ions (OsPIP2.4) (Ono *et al*., 2025; Qiu *et al*., 2025), CO_2_ (AtPIP2.1) (Chen *et al*., 2023), urea (AtTIP5.1) (Dynowski *et al*., 2008*a*), NH_3_ (AtTIP2.1) (Kirscht *et al*., 2016), boron (AtNIP5.1), silicon (OsNIP2.1) (Mitani-Ueno *et al*., 2011) and glycerol (LIMP2) (Supplementary Fig. S5–S8). For each isoform, we examined the NPA motifs (loops B and E), the ar/R filter positions (H2, H5, LE1, LE2), and Froger’s positions (P1–P5). We then performed an *in silico* analysis to infer the putative substrates transported by each isoform.

**Table 2.** Conserved selectivity-related residues and predicted substrate permeability of Allium cepa aquaporins.

| Gene ID | Gene name | NPA<sup>a</sup> motif<br>LB/LE<sup>a</sup> | ar/R selectivity filter |  |  |  | Froger's positions |  |  |  |  | non-aqua putative transport |
| --- | --- | --- | --- | --- | --- | --- | --- | --- | --- | --- | --- | --- |
|  |  |  | H2 | H5 | LE1 | LE2 | P1 | P2 | P3 | P4 | P5 |  |
| g471398.t1 | AcPIP2.1 | NPA/NPA | F | H | T | R | Q | S | A | F | W | CO<sub>2</sub><sup>abcln</sup>, H<sub>2</sub>O<sub>2</sub><sup>abn</sup>, B<sup>*1</sup>, Urea<sup>*1</sup>, Ion<sup>n*2</sup> |
| g269457.t1 | AcPIP2.2 | NPA/NPA | F | H | T | R | Q | S | A | F | W | CO<sub>2</sub><sup>abcln</sup>, H<sub>2</sub>O<sub>2</sub><sup>abn</sup>, B<sup>*1</sup>, Urea<sup>*1</sup>, Ion<sup>n*2</sup> |
| g163028.t1 | AcPIP2.3 | NPA/NPA | F | H | T | R | Q | S | A | F | W | CO<sub>2</sub><sup>abcln</sup>, H<sub>2</sub>O<sub>2</sub><sup>abn</sup>, B<sup>*1</sup>, Urea<sup>*1</sup>, Ion<sup>n*2</sup> |
| g510823.t1 | AcPIP2.4 | NPA/NPA | F | H | T | R | Q | S | A | F | W | CO<sub>2</sub><sup>abcln</sup>, H<sub>2</sub>O<sub>2</sub><sup>abn</sup>, B<sup>*1</sup>, Urea<sup>*1</sup>, Ion<sup>n*2</sup> |
| g192632.t1 | AcPIP2.5 | NPA/NPA | F | H | T | R | R | S | A | F | W | CO<sub>2</sub><sup>l</sup>, Ion<sup>m</sup> |
| g454913.t1 | AcPIP1.1 | NPA/NPA | F | H | T | R | Q | S | A | F | W | CO<sub>2</sub><sup>abcln</sup>, H<sub>2</sub>O<sub>2</sub><sup>abn</sup>, B<sup>*1</sup>, Urea<sup>*1</sup>, Ion<sup>*2</sup> |
| g345607.t1 | AcPIP1.2 | NPA/NPA | F | H | T | R | Q | S | A | F | W | CO<sub>2</sub><sup>abcln</sup>, H<sub>2</sub>O<sub>2</sub><sup>abn</sup>, B<sup>*1</sup>, Urea<sup>*1</sup>, Ion<sup>*2</sup> |
| g103868.t1 | AcPIP1.3 | NPA/NPA | V | H | T | R | E | S | A | F | W | CO<sub>2</sub><sup>l</sup> |
| g259892.t1 | AcPIP1.4 | NPA/NPA | V | H | T | R | Q | S | A | E | K | - |
| g333327.t1 | AcTIP1.1 | NPA/NPA | H | I | A | V | T | S | A | Y | W | H<sub>2</sub>O<sub>2</sub><sup>a*1*2*3</sup>, NH<sub>3</sub><sup>ajh*1*2*3</sup>, Urea<sup>abd*1*2*3</sup>, B<sup>a*1*3</sup>, Gly<sup>a*3</sup> |
| Ace7g519610 | AcTIP1.2 | NPA/NPA | H | I | A | V | T | A | A | Y | W | CO<sub>2</sub><sup>l</sup>, H<sub>2</sub>O<sub>2</sub><sup>a*2</sup>, NH<sub>3</sub><sup>ajh*2</sup>, Urea<sup>abd*2</sup>, B<sup>a*1</sup>, Gly<sup>a</sup> |
| g455385.t1 | AcTIP1.3 | NPA/NPA | H | I | A | V | T | A | A | Y | W | CO<sub>2</sub><sup>l</sup>, H<sub>2</sub>O<sub>2</sub><sup>a*2</sup>, NH<sub>3</sub><sup>ajh*2</sup>, Urea<sup>abd*2</sup>, B<sup>a*1</sup>, Gly<sup>a</sup> |
| g163470.t1 | AcTIP2.1 | NPA/NPA | H | I | G | R | T | S | A | Y | W | H<sub>2</sub>O<sub>2</sub><sup>c*2</sup>, NH<sub>3</sub><sup>abchj*2</sup>, Urea<sup>bcd</sup>, Gly<sup>*2</sup> |
| g163473.t1 | AcTIP2.2 | NPA/NPA | H | I | G | R | T | S | A | Y | W | H<sub>2</sub>O<sub>2</sub><sup>c*2</sup>, NH<sub>3</sub><sup>abchj*2</sup>, Urea<sup>bcd</sup>, Gly<sup>*2</sup> |
| g163471.t1 | AcTIP2.3 | NPA/NPA | H | I | G | R | T | S | A | Y | W | H<sub>2</sub>O<sub>2</sub><sup>c*2</sup>, NH<sub>3</sub><sup>abchj*2</sup>, Urea<sup>bcd</sup>, Gly<sup>*2</sup> |
| g494803.t1 | AcTIP2.4 | NPA/NPA | H | I | G | R | T | S | A | Y | W | H<sub>2</sub>O<sub>2</sub><sup>c*2</sup>, NH<sub>3</sub><sup>abchj*2</sup>, Urea<sup>bcd</sup>, Gly<sup>*2</sup> |
| g494804.t1 | AcTIP2.5 | NPA/NPA | H | I | G | R | T | S | A | Y | W | H<sub>2</sub>O<sub>2</sub><sup>c*2</sup>, NH<sub>3</sub><sup>abchj*2</sup>, Urea<sup>bcd</sup>, Gly<sup>*2</sup> |
| g290563.t1 | AcTIP2.6 | NPA/NPA | H | I | G | R | T | S | A | Y | W | H<sub>2</sub>O<sub>2</sub><sup>c*2</sup>, NH<sub>3</sub><sup>abchj*2</sup>, Urea<sup>bcd</sup>, Gly<sup>*2</sup> |
| g41605.t1 | AcTIP2.7 | NPA/NPA | H | I | G | R | T | S | A | Y | W | H<sub>2</sub>O<sub>2</sub><sup>c*2</sup>, NH<sub>3</sub><sup>abchj*2</sup>, Urea<sup>bcd</sup>, Gly<sup>*2</sup> |
| g290561.t1 | AcTIP2.8 | NPA/NPA | H | I | G | R | T | S | A | Y | W | H<sub>2</sub>O<sub>2</sub><sup>c*2</sup>, NH<sub>3</sub><sup>abchj*2</sup>, Urea<sup>bcd</sup>, Gly<sup>*2</sup> |
| g290555.t1 | AcTIP2.9 | NPA/NPA | H | I | G | R | T | S | A | Y | W | H<sub>2</sub>O<sub>2</sub><sup>c*2</sup>, NH<sub>3</sub><sup>abchj*2</sup>, Urea<sup>bcd</sup>, Gly<sup>*2</sup> |
| g362212.t1 | AcTIP2.10 | NPA/NPA | H | I | G | R | T | S | A | Y | - | H<sub>2</sub>O<sub>2</sub><sup>c*2</sup>, NH<sub>3</sub><sup>abchj*2</sup>, Urea<sup>bcd</sup>, Gly<sup>*2</sup> |
| g330520.t1 | AcTIP3.1 | NPA/NPA | H | I | A | R | T | A | A | Y | W | H<sub>2</sub>O<sub>2</sub><sup>*2</sup>NH<sub>3</sub><sup>ah</sup>, Urea<sup>*2</sup> |
| Ace3g237400 | AcTIP3.2 | NPA/NPA | H | I | A | R | T | A | A | Y | W | H<sub>2</sub>O<sub>2</sub><sup>*2</sup>NH<sub>3</sub><sup>ah</sup>, Urea<sup>*2</sup> |
| g406039.t1 | AcTIP4.1 | NPA/NPA | H | I | A | R | T | S | A | Y | W | NH<sub>3</sub><sup>abh</sup>, Urea<sup>bd*2</sup> |
| g520887.t1 | AcTIP5.1 | NPA/NPA | Q | I | A | R | S | S | A | Y | W | - |
| g342104.t1 | AcNIP1.1 | NPA/NPA | W | V | A | R | F | T | A | Y | I | H<sub>2</sub>O<sub>2</sub><sup>g</sup>, Gly<sup>e</sup> |
| g171711.t1 | AcNIP3.1 | NPA/NPA | W | V | A | R | F | S | A | Y | I | H<sub>2</sub>O<sub>2</sub><sup>ag*2</sup>, As<sup>c*2*3</sup>, Sb<sup>*2</sup>, B<sup>*1</sup>, NH<sub>3</sub><sup>b*1</sup>, Gly<sup>ce*1*2</sup> |
| g171774.t1 | AcNIP4.1 | NPA/NPA | W | V | A | R | L | S | A | Y | I | H<sub>2</sub>O<sub>2</sub><sup>g</sup>, NH<sub>3</sub><sup>b</sup>, Gly<sup>e</sup> |
| g525889.t1 | AcNIP2.1 | NPA/NPA | G | S | A | R | M | T | A | Y | F | Si<sup>bik</sup>, B<sup>b</sup>, Urea<sup>b</sup> |
| g106325.t1 | AcNIP2.2 | NPA/NPA | G | A | G | R | F | T | A | Y | F | - |
| g428538.t1 | AcNIP5.1 | NPS/NPV | S | I | A | R | F | T | A | Y | L | B<sup>k</sup> |
| g368937.t1 | AcNIP5.2 | NPS/NPV | A | I | A | R | F | T | A | Y | I | As<sup>a*2</sup>, B<sup>bk*2</sup>, Urea<sup>af*2</sup>, Gly<sup>af*2</sup> |
| g64935.t1 | AcNIP7.1 | NPA/NPA | W | G | F | R | Y | D | K | Y | F | As<sup>o</sup>, Sb<sup>o</sup>, Gly<sup>eo</sup> |
| g418330.t1 | AcSIP1.1 | NPV/NPA | I | V | P | N | M | A | A | Y | W | - |
| g32459.t1 | AcSIP1.2 | NPA/NPA | I | I | P | N | M | A | A | Y | W | - |
| g146897.t1 | AcSIP2.1 | NPL/NPA | F | H | G | S | F | A | A | Y | W | - |
| Ace8g588110 | AcSIP2.2 | NPL/NPA | S | H | G | S | F | A | A | Y | W | - |
| g204256.t1 | AcSIP2.3 | NPL/NPA | S | H | G | A | F | A | A | Y | W | - |
| g145045.t1 | AcSIP2.4 | NPL/NPA | S | H | G | S | F | A | A | Y | W | - |

The conserved NPA/NPA motif in loops B and E was found in all PIP and TIP isoforms, in six of the eight NIP isoforms, and in AcSIP1.2. AcNIP5.1 and AcNIP5.2 displayed the variants NPS/NPV. Among the SIPs, AcSIP1.1 showed NPV/NPA, whereas all SIP2 isoforms carried NPL in loop B.

Nearly all PIP sequences carried the canonical residues associated with water transport (Otto *et al*., 2010) in both the ar/R filter (F, H, T, R) and Froger’s positions P2–P4 (S, A, F, W) (Table 2). Phylogenetic analysis also allowed us to expand our substrate prediction to include CO_2_ and H_2_O_2_ (Hove and Bhave, 2011; Perez Di Giorgio *et al*., 2014; Azad *et al*., 2016). We identified three notable deviations from the canonical water-transporter pore: AcPIP2.5 displayed an arginine (R) at P1 instead of the usual glutamine (Q), AcPIP1.3 carried glutamate (E) at P1 and valine (V) at the H_2_ position (Supplementary Fig. S9), and AcPIP1.4 also had valine at H_2_, plus additional residues inserted in helix 6 and substitutions at P4 (F→E) and P5 (W→K) (Supplementary Fig. S10). Because the combination of novel substitutions in AcPIP1.4 has not been observed in other plant aquaporins, we were unable to predict its non-water substrate permeability. However, AcPIP1.3 contains additional amino-acid residues outside the canonical filter and Froger’s positions that match those previously shown to permit CO_2_ transport (Chen *et al*., 2023). Similarly, AcPIP2.5 not only retains the CO_2_-permeability signature but also displays features characteristic of ion-conducting aquaporins (Ono *et al*., 2025) (Supplementary Fig. S5).

All AcTIP sequences possessed the conserved ar/R filter and Froger’s signature characteristic of tonoplast channels, consistent with their primary roles in water and nitrogen transport. In silico predictions indicated that both TIP1 (group I) and TIP2 (group IIa) isoforms can facilitate NH_3_ and urea transport. Literature reports have additionally linked the TIP1 subgroup to boron and glycerol permeability, whereas TIP2 isoforms lack the classical determinants for these larger solutes (Azad *et al*., 2012). However, comparative homology with known glycerol transporters in model plants suggests that glycerol may also be a putative substrate for TIP2 isoforms. Novel insights into CO_2_ transport-related residues also suggest that some TIP1 isoforms can facilitate CO_2_ permeability (Supplementary Fig. S6). Group IIb (TIP3 and TIP4) (Azad *et al*., 2016) isoforms were predicted to transport NH_3_, and AcTIP4.1 was additionally predicted to transport urea (Hove and Bhave, 2011). Potential H_2_O_2_ transport by TIPs cannot be ruled out based on homology with Arabidopsis TIP aquaporins (Perez Di Giorgio *et al*., 2014). AcTIP5.1 was the only isoform exhibiting high variability in both the ar/R filter (Q, I, A, R) and Froger’s positions (S, S, A, Y, W) (Supplementary Fig. S6 and Supplementary Fig. S11). However, this level of variability is typical for TIP5; similar patterns have been observed in OsTIP5.1 (rice) and ZmTIP5.1 (maize), but no experimental analysis has yet determined the actual substrates transported by TIPs5. Consequently, no reliable substrate prediction could be made for AcTIP5.1 (Table 2).

AcNIP1.1, AcNIP3.1, and AcNIP4.1 all share the classical glycerol-transporter ar/R signature (W, V, A, R). Based on previous studies, these three isoforms were also predicted to transport H_2_O_2_ (Dynowski *et al*., 2008*b*). Using homology models (Perez Di Giorgio *et al*., 2014), AcNIP3.1 is additionally predicted to transport arsenic, boron, and NH_3_ alongside glycerol and H_2_O_2_, whereas AcNIP4.1 was predicted only for NH_3_. AcNIP7.1 exhibits the same Ar/R signature as the prototypical glycerol transporter from Escherichia coli (GlpF) which is capable of transporting glycerol and other polyols. (Pluhackova *et al*., 2022). AcNIP2.1 grouped with known silicon transporters such as OsNIP2.1 (Fig. 1), and its ar/R and Froger’s residues matched those required for Si permeability (Mitani-Ueno *et al*., 2011). Additionally, potential boron and urea transport could not be ruled out (Hove and Bhave, 2011). Although AcNIP2.2 clusters adjacent to AcNIP2.1 in the phylogeny, the presence of an unusual glycine at LE_1_ precluded any reliable substrate prediction (Supplementary Fig. S7 and Supplementary Fig. S12). AcNIP5.1 and AcNIP5.2 share nearly identical residue signatures, except that AcNIP5.1 carries a serine (S) at the H_2_ position and G at the LE_1_. Consequently, AcNIP5.2 is predicted to transport multiple solutes —including arsenic, boron, urea, and glycerol— whereas AcNIP5.1 is predicted only for boron (Mitani-Ueno *et al*., 2011) (Supplementary Fig. S7 and Supplementary Fig. S13). No substrate predictions could be made for AcSIP isoforms based on homology, orthology, or experimental studies (Supplementary Fig. S8).

### 3.5 Physiological and biochemical data

Transpiration rate, stomatal conductance (gs) and net photosynthesis were significantly higher in set-derived plants compared with seed-derived ones (Fig. 2A, 2B and 2C respectively). In contrast, the intracellular CO_2_ concentration (Ci) was significantly lower in set-derived plants (Fig. 2D). Photosynthetic CO_2_ assimilation (A/PAR) also showed a marked increase in set-derived plants relative to seed-derived ones (Fig. 2E). No significant differences were detected between propagation types for water use efficiency (WUE) or apparent carboxylation efficiency (ACE) (Fig. 2F and 2G), indicating that the intrinsic biochemical capacity for CO_2_ fixation was comparable in both propagation types. No significant differences were observed in total plant weight (Fig. 2H).

**Fig. 2.**
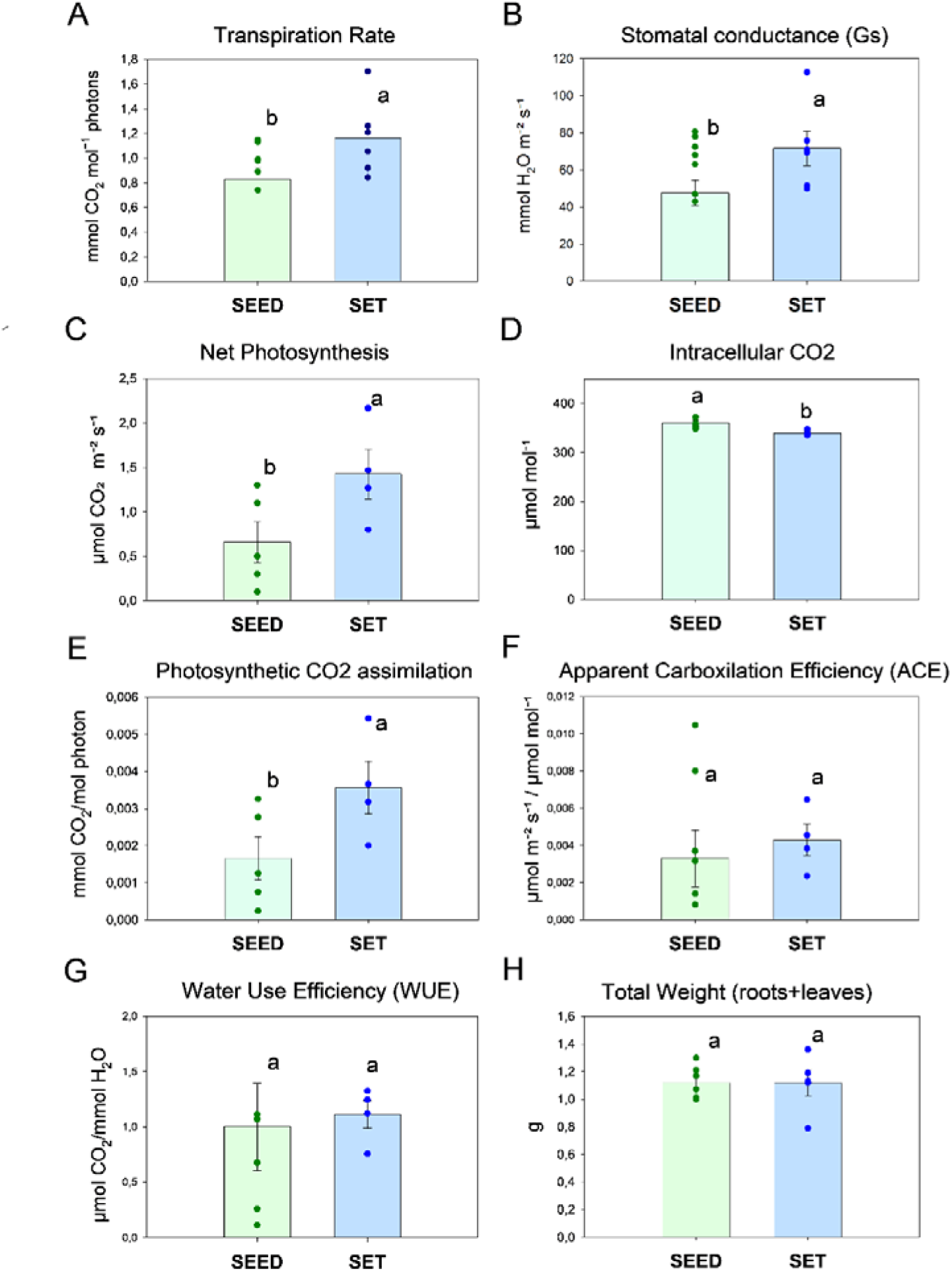
Physiological parameters measured in onion plants derived from seed (SEED, green) or onion set (SET, blue). **(A)** Transpiration rate, **(B)** stomatal conductance (Gs), **(C)** net photosynthesis, **(D)** intracellular CO_2_ concentration (Ci), **(E)** photosynthetic CO_2_ assimilation rate (A/PAR), **(F)** apparent carboxylation efficiency (ACE), **(G)** instantaneous water use efficiency (WUE) and **(H)** total weight (including roots and leaves). Bars represent mean values ± standard error (n = 10). Individual dots represent biological replicates. Differences between propagation modes were analysed by one-way ANOVA followed by Duncan’s multiple range post hoc test (p < 0.05). Different letters above bars indicate statistically significant differences **Alt text:** Multi-panel bar chart (A–H) comparing physiological traits between seed-derived and set-derived onion plants. Each panel shows two bars representing the two propagation types, with individual replicate points and error bars. Variables include transpiration, stomatal conductance, photosynthesis, intercellular carbon dioxide concentration, photosynthetic efficiency, carboxylation efficiency, water use efficiency, and total fresh weight. Letters above bars indicate statistical groupings.

Seed-derived plants consistently maintained high hydrogen peroxide (H_2_O_2_) levels in all tissues analysed (leaf, root and bulb) while set-derived plants exhibited significantly lower H_2_O_2_ accumulation in leaves and roots, with a marked increase only in the bulb (Fig. 3A). Regarding boron content, seed-derived plants showed a more polarized distribution, with high levels in leaves and notably lower concentrations in roots and bulbs. In contrast, set-derived plants displayed a more homogeneous boron profile across tissues, with moderate to high levels in all organs and significantly higher values in root and bulb compared to seed-derived plants (Fig. 3B).

**Fig. 3.**
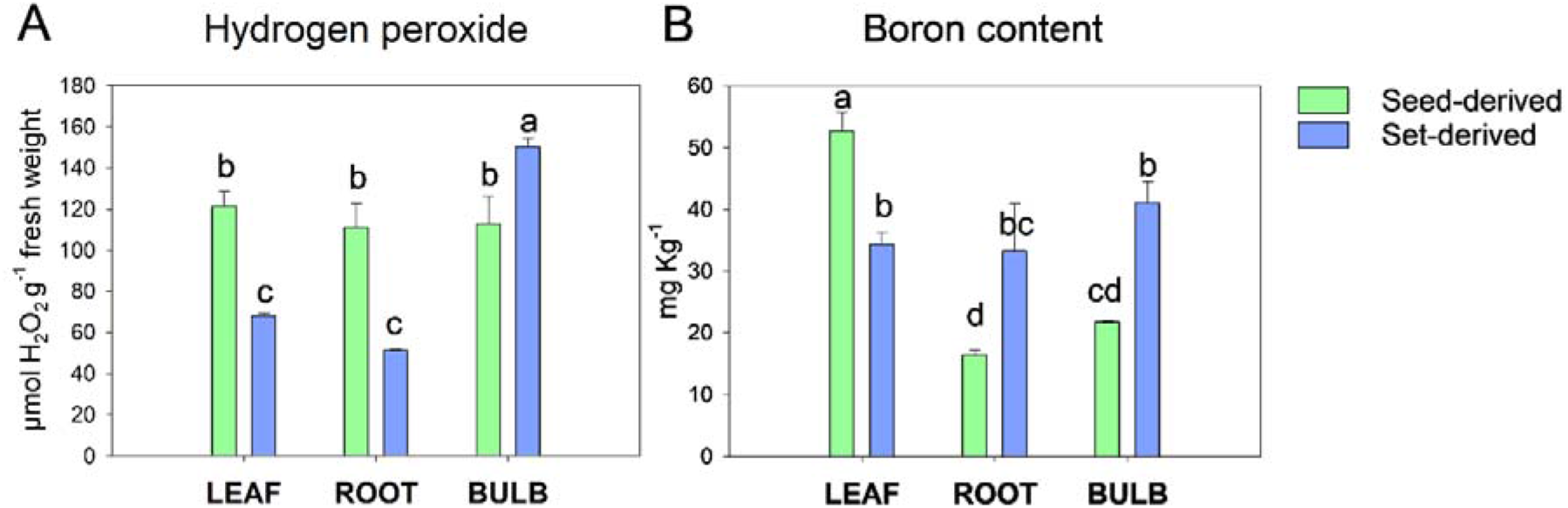
Biochemical differences between seed-derived (Green) and set-derived (Blue) onion plants in leaf, root and bulb tissues. **(A)** Hydrogen peroxide (H_2_O_2_) content and **(B)** Boron concentration. Bars represent mean values ± standard error (n = 6 and n = 3 biological replicates per propagation mode and tissue respectively). Differences between propagation modes were analysed by one-way ANOVA followed by Duncan’s multiple range post hoc test (p < 0.05). Different letters above bars indicate statistically significant differences. Individual biological replicate values are provided in Supplementary Table S1. **Alt text:** Two bar charts comparing biochemical parameters in leaf, root, and bulb tissues of seed-derived and set-derived onion plants. Panel A shows hydrogen peroxide content and panel B shows boron concentration. Bars are colour-coded by propagation type, include error bars, and letters above bars indicate statistical groupings.

### 3.6 Expression analysis of AcAQPs

We compiled RNA-seq data from various BioProjects (Supplementary Table S3) and organized it into a stacked bar chart graph and heatmaps showing AcAQP expression changes across different tissues (Fig. 4). The expression patterns of the *AcAQP* genes were examined across six onion tissues (leaf, leaf sheath, bulb, root, floral bud and flower) using RNA-seq data normalized by the VST method. Constitutive expression varied markedly among isoforms (Fig. 4A). The most highly expressed aquaporins across tissues belonged mainly to the PIP and TIP subfamilies, particularly *AcPIP1*.*1, AcPIP1*.*2, AcPIP2*.*4, AcTIP2*.*7* and *AcTIP1*.*1*. Within the NIP subfamily, *AcNIP5*.*2* stood out as the most expressed isoform, while among SIPs, *AcSIP1*.*1* and *AcSIP1*.*2* were the most abundant members.

**Fig. 4.**
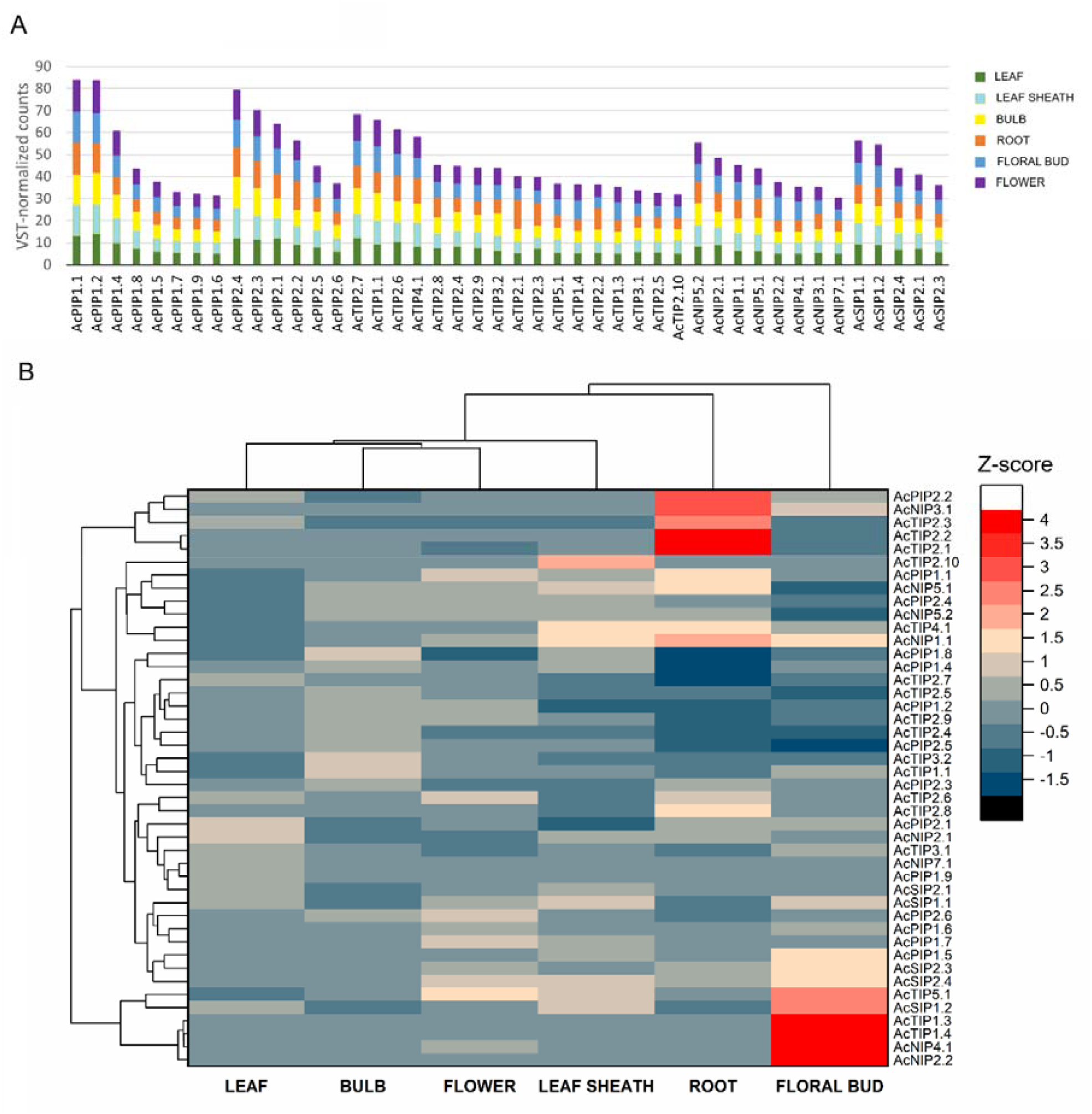
Constitutive expression profiles of aquaporin genes across onion tissues based on public RNA-seq datasets. **(A)** Stacked bar chart showing mean VST-normalized expression values (DESeq2) of each aquaporin gene across six onion tissues: leaf (LE), leaf sheath (LS), bulb (BU), root (RO), floral bud (FB) and flower (FL). Genes are grouped by subfamily (PIPs, TIPs, NIPs and SIPs) and ordered by decreasing overall expression. **(B)** Tissue-specific expression heatmap of AcAQPs. Hierarchical clustering of 48 AcAQP isoforms based on VST-normalized RNA-seq counts across six onion tissues (leaf, bulb, flower, leaf sheath, root, floral bud). Each cell’s colour represents the relative expression of a given isoform per tissue (red = highest; dark blue = lowest). Dendrograms group aquaporin isoforms (rows) and tissues (columns) according to expression similarity. Data were compiled from multiple BioProjects (detailed in Supplementary Table S3). **Alt text:** Two-panel figure showing constitutive aquaporin gene expression across six onion tissues. Panel A is a stacked bar chart showing normalized expression values for each aquaporin gene, with colours representing different tissues. Panel B is a heatmap showing tissue-specific relative expression patterns of aquaporin genes, with rows representing genes, columns representing tissues, and colours indicating relative expression levels. Dendrograms show clustering based on expression similarity.

Heatmap analysis based on z-score normalization revealed distinct tissue-specific expression patterns among AcAQPs (Fig.4B). The most striking tissue-specific changes in aquaporins expression occurred in roots, where several isoforms from PIPs, TIPs and NIPs were highly represented, particularly *AcPIP1*.*1, AcPIP2*.*2*, multiple TIP2 isoforms, *AcNIP1*.*1, AcNIP3*.*1* and *AcNIP5*.*1*. The floral bud exhibited its own distinct expression cluster, with *AcPIP1*.*5* as sole highlighted PIP, *AcTIP1*.*3, AcTIP1*.*4, AcTIP5*.*1* as TIPs, NIPs (*AcNIP2*.*2, AcNIP4*.*1*), and almost all SIPs. In leaves and bulbs, subgroup-level shifts were less pronounced, although certain isoforms stood out. In leaves, *AcPIP2*.*1, AcTIP2*.*7* and *AcNIP2*.*1* were the most highly expressed genes, whereas bulbs preferentially expressed several PIP and TIP isoforms, particularly *AcPIP1*.*8, AcTIP1*.*1* and *AcTIP3*.*2*. Leaf sheaths were characterized by high *AcTIP2*.*10* expression together with elevated *AcTIP4*.*1, AcNIP1*.*1* and SIPs1 levels. Flowers showed increased expression of *AcPIP1*.*7, AcPIP2*.*6, AcTIP2*.*6, AcTIP5*.*1* and *AcNIP4*.*1* although the latter two reached their highest expression levels in floral buds.

Based on the experimental RNA-seq data, clear propagation-dependent expression patterns were observed in both roots and leaves (Fig. 5). In roots, set-derived plants showed higher expression of multiple PIP2 and TIP2 isoforms, together with *AcTIP4*.*1*, whereas seed-derived plants preferentially expressed several PIP1, NIP and SIP genes (Fig. 5), particularly some PIPs1 aquaporins and specific NIPs (*AcNIP1*.*1, AcNIP2*.*1, AcNIP3*.*1, AcNIP5*.*1* and *AcNIP5*.*2*). A similar pattern was observed in leaves, where set-derived plants displayed enhanced expression of several TIP2 and PIP isoforms and als*AcTIP1*.*1* was also highly expressed in set-derived leaves. In contrast, seed-derived leaves showed higher expression of selected PIP1s, *AcPIP2*.*6* as the only PIP2, some TIP (*AcTIP1*.*4, AcTIP3*.*1* and *AcTIP5*.*1), AcNIP2*.*1* and SIP isoforms.

**Fig. 5.**
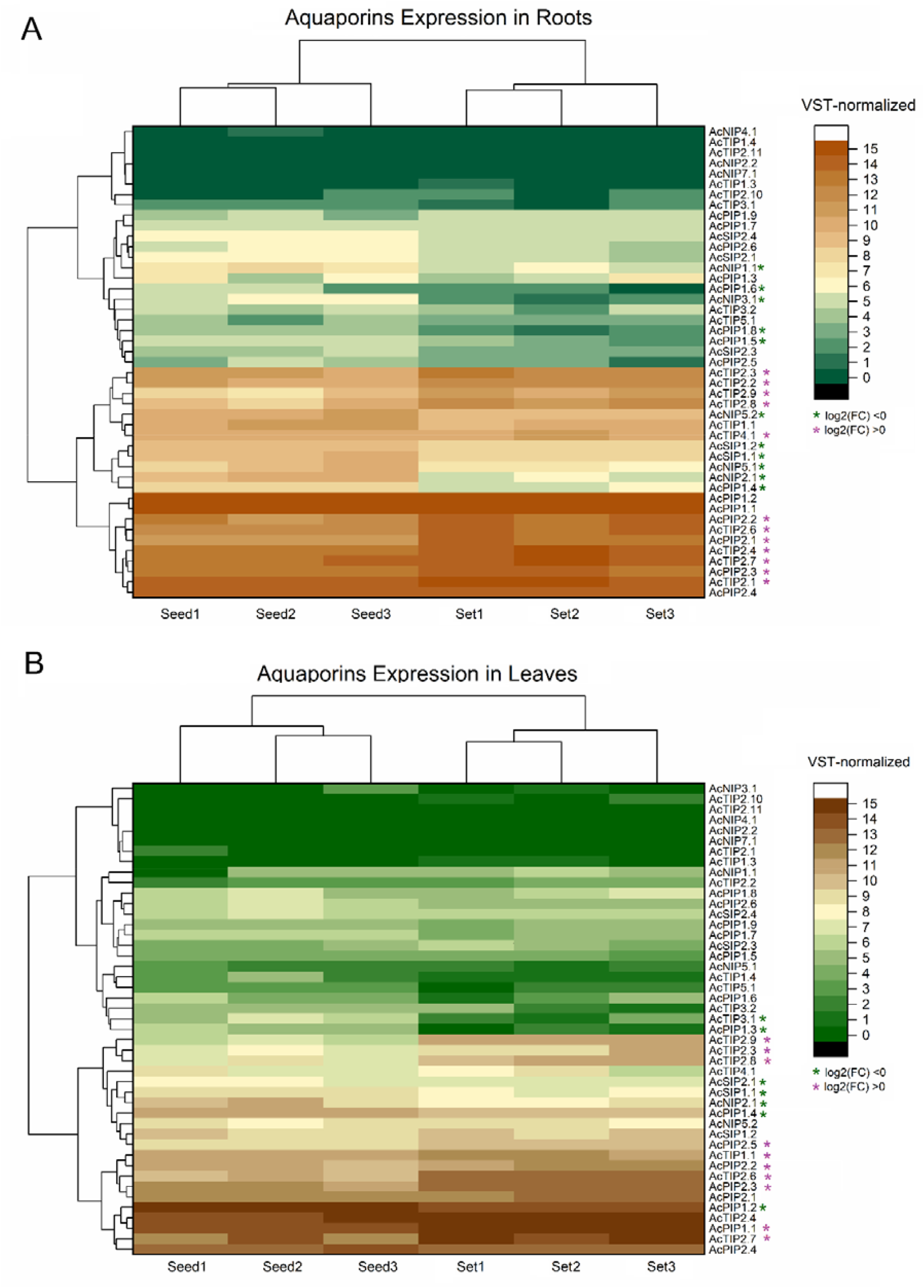
Aquaporin gene expression patterns in (A) roots and (B) leaves of seed-derived and set-derived onion plants. Heatmaps show VST-normalized expression values (DESeq2) for 48 AcAQP genes in three biological replicates per propagation mode and tissue (Seed1–3 and Set1–3). Colours indicate increasing normalized expression levels according to the scale shown. Dendrograms represent hierarchical clustering of samples and genes based on expression similarity. Asterisks indicate genes showing significant differential expression between propagation modes according to DESeq2 (adjusted p-value ≤ 0.05 and |log_2_ fold change| ≥ 0.5). Green asterisks indicate genes upregulated in seed-derived plants (log_2_FC < 0), whereas purple asterisks indicate genes upregulated in set-derived plants (log_2_FC > 0). **Alt text:** Two heatmaps showing aquaporin gene expression in roots (top) and leaves (bottom) of seed-derived and set-derived onion plants. Columns represent biological replicates from each propagation type and rows represent aquaporin genes. Colours indicate normalized expression levels according to the scale shown. Dendrograms indicate clustering of samples and genes based on expression similarity. Coloured asterisks mark genes with significant differential expression between propagation types.

## 4. Discussion

Our results demonstrate that propagation mode can shape whole-plant growth strategy through the differential deployment of aquaporin transport networks in onion. These contrasting strategies are supported by differences in both the composition and expression patterns of the aquaporin gene family.

The aquaporin gene family comprises 48 members in *Allium cepa* (AcAQPs), including 39 full-length sequences and 9 partial sequences. While some sequences appear incomplete, their detectable expression in transcriptomic datasets suggests that assembly gaps—common in the highly repetitive onion genome—are the most likely explanation (Finkers *et al*., 2021). Phylogenetic analysis grouped AcAQPs into the major plant aquaporin subfamilies (PIPs, TIPs, NIPs and SIPs), with a distribution comparable to that reported in other plant species (Supplementary Table S5). Notably, the relatively high representation of PIP1 and TIP2 isoforms may be functionally relevant, given their established roles in water and solute transport. Two isoforms, AcPIP1.1 and AcPIP1.2, clustered with PIP1 homologs from other taxa (Fig. 1), consistent with an ancient duplication possibly retained across monocots (Zou and Yang, 2019). In contrast, the remaining PIP1 isoforms and TIP2 members formed onion-specific clusters and were located in close proximity within the genome (Supplementary Table S4), suggesting more recent tandem or segmental duplication events that aligns with functional specialization under lineage-specific selection pressure (Johanson *et al*., 2001; Nicolas-Espinosa and Carvajal, 2022). Such localised expansions are common in transport-related gene families, where gene dosage and expression flexibility may confer adaptive advantages and functional specialization (Hanada *et al*., 2008). Taken together, the expansion patterns observed in both PIP1 and TIP2 subfamilies reflect a combination of ancestral and more recent duplication events.

The biochemical and structural analyses further support a conserved specialization of aquaporin subfamilies across species, consistent with patterns observed in other plant systems (Lopez-Zaplana *et al*., 2020; Malik *et al*., 2023). Predicted subcellular localisation matched expectations for most isoforms, with a few exceptions that will require experimental confirmation. Solute transport predictions were carried out using the same structural criteria established by Lopez-Zaplana et al. 2020 (Lopez-Zaplana *et al*., 2020), updated with more recent literature identifying key residues involved in the transport of CO_2_, H_2_O_2_, Si and ions (Van den Berg *et al*., 2021; Chen *et al*., 2023; Ono *et al*., 2025; Qiu *et al*., 2025). For isoforms presenting novel or unusual motifs within the Ar/R selectivity filter or Froger’s positions, three-dimensional structure modelling was performed to assess potential substrate preferences. Key structural insights are discussed in Supplementary Fig S8–S14. Together, these analyses support the idea that even within conserved subfamilies, specific isoforms may have evolved distinct solute affinities or membrane localisation patterns, contributing to the functional diversity observed in onion AQPs.

When studying the response of both seed- and set-derived plants at the same developmental stage (similar weight, Fig. 2H), set-derived plants reached the same developmental stage substantially faster despite being planted 47 days later than seed-derived plants. This temporal advantage highlights the accelerated post-dormancy growth capacity of sets. Accordingly, physiological parameters in set-derived plants displayed significantly higher stomatal conductance (Gs), transpiration rate (E) and net photosynthesis (A), alongside a lower intercellular CO_2_ concentration (Ci) (Fig. 2A–D). This pattern reflects a more active gas exchange and photosynthetic metabolism, where CO_2_ is rapidly assimilated once diffused into the leaf (Gobu *et al*., 2022). In contrast, seed-derived plants exhibited lower A and Gs values but higher Ci, indicative of more limited internal CO_2_ diffusion and a more conservative carbon-water exchange strategy. Despite these differences, neither apparent carboxylation efficiency (ACE = A/Ci) nor instantaneous water use efficiency (WUE = A/E) differed significantly between propagation types (Fig. 2F–G). This apparent similarity does not indicate the absence of physiological divergence, but rather reflects proportional changes in the underlying fluxes (Tshikunde *et al*., 2018; Guerrieri *et al*., 2019).

The hydrogen peroxide accumulation profile reveals a marked contrast in redox dynamics between the two propagation types. Seed-derived plants maintained elevated H_2_O_2_ levels across all analysed tissues, suggesting altered basal redox homeostasis and/or differences in ROS turnover capacity. Such constitutively higher ROS levels have been associated with a state of heightened physiological alertness and sustained engagement of antioxidant defence pathways, and are often linked to slower developmental progression even in the absence of visible stress symptoms (Smirnoff and Arnaud, 2019).

The distribution of boron also differed markedly between propagation types, consistent with differences in their underlying molecular transport systems. In seed-derived plants, the high boron concentration in leaves-despite lower root levels-points to preferential root-to-shoot allocation or remobilization. Such partitioning may enhance boron supply to aerial tissues, where this element plays essential roles in cell wall stabilization and developmental signalling (Matthes *et al*., 2020).

PIP aquaporins are major determinants of plasma membrane water permeability and whole-plant hydraulic transport in plants (Chaumont and Tyerman, 2014). *AcPIP1*.*1, AcPIP1*.*2* and *AcPIP2*.*4* were the most constitutively expressed aquaporins across onion tissues (Fig. 4A), pointing to central roles in basal hydraulic homeostasis. Notably, none of these three isoforms changed in roots of set- or seed-derived plants (Fig. 5), suggesting that high root water uptake capacity is conserved regardless of propagation type, and reinforcing their central role in root hydraulics. In leaves, however, these PIP1 isoforms showed differential regulation between propagation methods, with a higher expression of *AcPIP1*.*1* in set-derived leaves while *AcPIP1*.*2* was more highly expressed in seed-derived plants. This contrasting expression pattern suggests alternative leaf hydraulic and carbon-acquisition strategies associated with each propagation system. Beyond water transport, several PIPs can facilitate CO_2_, ions or H_2_O_2_ movement, thereby influencing mesophyll conductance and photosynthetic performance (Bienert *et al*., 2007; Katsuhara and Hanba, 2008; Qiu *et al*., 2020), and have been associated with facilitating mesophyll CO_2_ diffusion to sustain photosynthesis (Flexas *et al*., 2006; Uehlein *et al*., 2012). Moreover, PIP1–PIP2 hetero-tetramerization increases PIP1 abundance at the plasma membrane and enhances water permeability (Bienert *et al*., 2018), while PIP1:PIP2 stoichiometry can tune relative CO_2_ vs. H_2_O conductance (Otto *et al*., 2010). As described previously, set-derived plants exhibited significantly higher stomatal conductance and transpiration rates compared to seed-derived plants (Fig. 2), which in turn translated into enhanced CO_2_ assimilation rates. The lower intercellular CO_2_ concentration (Ci) is consistent with increased CO_2_ consumption in the mesophyll, indicating a more active photosynthetic metabolism (Heckwolf *et al*., 2011). In addition, the apparent carboxylation efficiency (ACE) remained unchanged between groups, suggesting that the Rubisco carboxylation capacity and the biochemical machinery for CO_2_ fixation operate with similar intrinsic efficiency in both cases (Sharkey *et al*., 2007). Together, these results indicate that the higher net CO_2_ assimilation rate of set-derived plants was not due to higher carboxylation efficiency but rather to factors enhancing CO_2_ availability such as aquaporins. These is consistent with improved internal CO_2_ conductance or more efficient hydraulic supply potentially mediated by PIP aquaporins (Table 2). These data identify PIP aquaporins as likely coordinators of the carbon–water balance underlying contrasting propagation strategies.

TIP aquaporins are major tonoplast channels involved in vacuolar water and nitrogen transport (Loqué *et al*., 2005). TIPs may also contribute to vacuolar homeostasis by facilitating controlled movement of H_2_O_2_ and nitrogenous solutes to support redox signalling and metabolic adjustment (Katsuhara *et al*., 2008; Hachez and Chaumont, 2010). Heatmap analysis revealed that most TIP2 isoforms and several PIP genes were upregulated in set-derived plants relative to seed-derived plants in both roots and leaves. In roots, increased expression of these aquaporins likely contributes to enhanced hydraulic conductivity, promoting water uptake and transport to the shoot (Chaumont and Tyerman, 2014), supporting the higher transpiration demands observed in set-derived plants (Fig. 2). In leaves, elevated aquaporin expression may improve mesophyll water movement and facilitate coupling between water and CO_2_ transport, thereby helping to maintain stomatal opening and gas exchange under higher transpiration fluxes (Heinen *et al*., 2009). Together, these data support a scenario in which set-derived plants operate a higher-capacity hydraulic configuration capable of sustaining elevated stomatal conductance and CO_2_ assimilation without changes in carboxylation efficiency.

Accordingly, the increased expression of AcPIP1.1 in set-derived plants, together with abundant PIP2 isoforms such as AcPIP2.2, AcPIP2.3, AcPIP2.5 and AcPIP2.6 (Fig. 5), aligns with the physiological profile (A↑, Ci↓, Gs/Tr↑) and supports a cooperative role in water and CO_2_ transport. Conversely, the higher expression of *AcPIP1*.*2* in seed-derived leaves supports a more constitutive, homeostatic aquaporin network. In arabidopsis, AtPIP1;2 is one of the most stably expressed PIP1s under non-stress conditions (Postaire *et al*., 2010), and its expression pattern resembles the seed-derived profile (lower A and Gs, higher Ci), consistent with steadier water transport.

Expression profiling of PIPs across onion tissues revealed marked tissue-specific patterns consistent with specialized physiological roles (Fig. 4). The differential expression of PIP1 isoforms across tissues likely reflects their evolutionary diversification. Distinct tissues displayed preferential PIP1 isoforms (Fig. 4B): *AcPIP1*.*1* in roots; *AcPIP1*.*2, AcPIP1*.*4* and *AcPIP1*.*8* in bulbs; *AcPIP1*.*9* in leaves; *AcPIP1*.*4* in leaf sheath; and *AcPIP1*.*5* and *AcPIP1*.*7* in floral tissues. Such partitioning may help explain why PIP1 genes are more numerous than PIP2s in onion. This specialization likely enables fine regulation of tissue-specific water transport demands. AcPIP1.1 showed dominant root expression, suggesting a central role in radial water transport and root hydraulic conductivity, as also reported for PtoPIP1;1 in *Populus tomentosa* (Leng *et al*., 2021). *AcPIP1*.*4* was highly expressed in leaf sheath and bulbs (Fig. 4B) and was also upregulated in both roots and leaves of seed-derived plants (Fig. 5). This pattern, combined with its sequence features (Supplementary Fig. S10), suggests a role in solute balance or signalling under a more conservative physiological strategy. Seed-derived plants, which showed higher internal CO_2_ and lower photosynthetic rates, also accumulated more hydrogen peroxide in both tissues (Fig. 3A) consistent with a role in cellular homeostasis and ROS signalling rather than bulk water transport. AcPIP1.5 and AcPIP1.7 were preferentially expressed in floral tissues, suggesting roles in reproductive stages (Fig. 4B). This is consistent with reports in other species where PIP aquaporins have been involved in petal expansion in rose, tulip, lily, carnations or gentians, among others (Ding *et al*., 2004; Azad *et al*., 2008; Katsuhara *et al*., 2008; Ma *et al*., 2008; Ren *et al*., 2025). Together, these observations support functional specialization of PIP1 isoforms following gene expansion, enabling fine regulation of tissue-specific water transport demands.

Among PIP2s, AcPIP2.2 emerged as the dominant plasma membrane PIP2 isoform in roots (Fig. 4) and was particularly abundant in set-derived plants (Fig. 5), pointing to a key role in water uptake during active growth. *AcPIP2*.*4* remained highly expressed across tissues, consistent with a robust basal system for water mobilization. Beyond water transport, several PIP2 isoforms may also contribute to ROS homeostasis through H_2_O_2_ diffusion across membranes (Bienert and Chaumont, 2014; Chaumont and Tyerman, 2014). In this context, *AcPIP2*.*5* was upregulated in set-derived plants (Fig. 5) and its particular sequence features suggest potential H_2_O_2_ permeability (Table 2). Notably, hydrogen peroxide concentration was higher in leaves of set-derived plants (Fig. 3A), consistent with a potential role in ROS redistribution.

Altogether, the contrasting transcriptional behaviour of PIP isoforms indicates substantial functional specialization. Constitutively expressed members such as AcPIP1.1, AcPIP1.2 and AcPIP2.4 may sustain basal hydraulic continuity, whereas tissue- and condition-responsive isoforms (e.g., AcPIP1.4, AcPIP2.2 and AcPIP2.5) likely provide regulatory flexibility. The differential modulation of PIPs between seed- and set-derived plants therefore appears to reflect adaptive tuning of water and CO_2_ transport capacities in line with the observed physiological performance.

The NIPs pattern (Fig. 5) suggests enhanced boron transport capacity in seed-derived plants. NIP aquaporins are major plasma membrane channels involved in metalloid transport (Pommerrenig *et al*., 2015). *AcNIP3*.*1, AcNIP1*.*1* and *AcNIP5*.*1* were the dominant constitutively expressed isoforms in roots. Interestingly, AcNIP3.1 and AcNIP5.1 are both related to boron transport (Table 2). Root-to-shoot boron transport has been well characterised in other species, where NIP5 and NIP3 family members play central roles (Takano *et al*., 2006; Diehn *et al*., 2019). Their expression in onion roots and floral buds (Fig. 4) where boron is essential (Jiang *et al*., 2025), supports a putative role in boron uptake and redistribution. Additionally, *AcNIP1*.*1* was highly expressed in roots and reproductive tissues. Although structurally predicted to mainly transport H_2_O_2_ and glycerol (Table 2), functional evidence from ZmNIP1.1 and other NIP1 isoforms indicates that this subfamily can also facilitate boron, ammonium, and urea transport (Bárzana *et al*., 2014; Lopez-Zaplana *et al*., 2022). These three isoforms-*AcNIP1*.*1, AcNIP3*.*1* and *AcNIP5*.*1*-were more highly expressed in roots of seed-derived plants, together with *AcNIP5*.*2* and *AcNIP2*.*1*, both of which may also contribute to boron transport based on their structural features (Table 2, Supplementary Fig. S12-S14). If functionally active in boron transport, these NIPs could support an active root-to-shoot mobilization, consistent with the higher boron accumulation detected in leaves of seed-derived plants (Fig. 3B).

Beyond metalloid transport, several NIPs have also been implicated in hydrogen peroxide diffusion across membranes, linking nutrient fluxes with redox signalling. In arabidopsis, AtNIP1;2 and AtNIP5;1 facilitate H_2_O_2_ transport (Dynowski *et al*., 2008*b*), and AtNIP1;1 contributes to hydrogen-peroxide sensitivity and is permeable to H_2_O_2_, linking NIP function to whole-plant redox responses (Sadhukhan *et al*., 2017). In rice, OsNIP3;2 and OsNIP3;3 also transport H_2_O_2_ (Katsuhara *et al*., 2014). Such roles suggest that NIPs may coordinate metalloid transport with redox signalling to maintain cellular homeostasis. This interpretation is consistent with the physiological divergence observed between propagation types. Set-derived plants displayed higher A, Gs and E values and lower Ci, indicative of efficient gas exchange and internal CO_2_ use. Seed-derived plants, in contrast, displayed lower A and Gs, higher Ci, and elevated H_2_O_2_ levels (Fig. 2 and Fig. 3A), consistent with a more conservative photosynthetic state accompanied by higher oxidative load. In this context, the higher expression of several NIP isoforms in seed-derived plants (*AcNIP1*.*1, AcNIP3*.*1, AcNIP5*.*1, AcNIP5*.*2* or *AcNIP2*.*1*) aligns with a greater requirement for coordinated solute and H_2_O_2_ transport to sustain redox equilibrium and metabolic stability. Overall, differential NIP deployment reflects divergent physiological priorities: set-derived plants emphasize high-efficiency hydraulic and carbon fluxes, whereas seed-derived plants rely on tighter internal regulation supported by enhanced metalloid and redox transport capacity.

## 5. Conclusions

This study provides the first genome-wide characterisation of the aquaporin family in *Allium cepa* (AcAQPs) and reveals that differential AQP deployment underlies the contrasting physiological behaviour of seed- and set-derived plants. Set-derived plants showed higher expression of AcPIP1.1, several PIP2 isoforms (AcPIP2.2, AcPIP2.3, AcPIP2.5, AcPIP2.6) and TIP2 members consistent with enhanced water and CO_2_ transport, higher stomatal conductance, transpiration and photosynthetic rates. This configuration supports a hydraulically efficient system capable of sustaining rapid carbon gain without evidence of altered biochemical fixation capacity.

In contrast, seed-derived plants exhibited higher expression of AcPIP1.2, AcPIP1.4 and several NIP isoforms (AcNIP1.1, AcNIP3.1, AcNIP5.1, AcNIP5.2, AcNIP2.1). These patterns coincided with higher H_2_O_2_ levels and greater boron accumulation in leaves, supporting a more tightly regulated internal environment focused on solute redistribution and redox balance rather than high-flux photosynthetic performance.

Altogether, our findings indicate that contrasting onion propagation types rely on distinct aquaporin-mediated transport strategies: a hydraulically optimized network promotes the accelerated growth of set-derived plants, whereas seed-derived plants depend on a solute- and redox-oriented network that may favour metabolic resilience. These insights highlight the key role of AQPs in defining growth dynamics and offer a molecular framework for future studies on water and nutrient transport in *Allium* crops. Propagation mode therefore appears not merely as a developmental alternative, but as a determinant of whole-plant growth strategy in onion.

## Supplementary data

The following supplementary data are available at JXB online.

**Supplementary Figure S1**. Principal component analysis (PCA) of VST-normalized expression data for aquaporin genes across six onion tissues.

**Supplementary Figure S2**. 3D protein structure prediction for A. cepa aquaporins.

**Supplementary Figure S3**. Nucleotide motif’s locations and sequences.

**Supplementary Figure S4**. Exon-Intron structure of the onion aquaporins genes.

**Supplementary Figure S5**. Multiple sequence alignment of onion PIPs with representative aquaporins characterised by the transport of water, CO_2_ and cations.

**Supplementary Figure S6**. Multiple sequence alignment of onion TIPs with representative aquaporins characterised by the transport of water, Urea and ammonia.

**Supplementary Figure S7**. Multiple sequence alignment of onion NIPs with representative aquaporins characterised by the transport of water, boron, silicon and glycerol.

**Supplementary Figure S8**. Multiple sequence alignment of onion SIPs with representative aquaporins characterised by the transport of water.

**Supplementary Figure S9**. Molecular representation of the onion aquaporins AcPIP1.2 and AcPIP1.3

**Supplementary Figure S10**. Predicted 3D structure model of AcPIP1.4

**Supplementary Figure S11**. Three-dimensional views of the Ar/R selectivity filter in AcTIP4.1 and AcTIP5.1.

**Supplementary Figure S12**. Three-dimensional views of the Ar/R selectivity filter in AcNIP2.1 and AcNIP2.2.

**Supplementary Figure S13**. Three-dimensional views of the Ar/R selectivity filter in AcNIP5.1 and AcNIP5.2.

**Supplementary Figure S14**. Three-dimensional views of the Ar/R selectivity filter in AcNIP3.1 and AcNIP7.1.

**Supplementary Table S1**. Individual biological replicate values used for statistical analyses and graphical representation of biochemical parameters.

**Supplementary Table S2**. Physicochemical properties of the identified A. cepa aquaporin proteins, including theoretical isoelectric point (pI), instability index, aliphatic index and GRAVY values.

**Supplementary Table S3**. Summary of RNA-seq datasets retrieved from the NCBI database and used for aquaporin expression analyses.

**Supplementary Table S4**. Chromosomal distribution of *Allium cepa* L. aquaporins.

**Supplementary Table S5**. Comparative distribution of aquaporin isoforms by subfamily in selected plant species, including model species, representative cereals and horticultural crops.

## Acknowledgements

Thanks to Dr. Yue Liu and Qingdao Agricultural University for their technical support in accessing the AlliumDB resources and website and Dr. Angel Luigi Guarnizo Serrudo for his technical advice.

## Author Contribution

Gloria Bárzana: Conceptualization, methodology, data curation, formal analysis, investigation, visualization, validation, writing, reviewing and editing, funding acquisition. Micaela Carvajal: Conceptualization, resources, validation, reviewing, supervision, funding.

## Conflict of interest

On behalf of all authors, the corresponding author states that there is no conflict of interest.

## Funding

This study was funded by the ‘Centro para el Desarrollo Tecnológico y la Innovación’ (CDTI) (Project number 20236980) supported by Ministerio de Ciencia, Innovación y Universidades (MCIN) and co-financed by Grupo Javaloyes S.L.

## Data availability

All aquaporin identifiers, curated annotations, predicted sequences and associated genome IDs used in this study are provided in Table 1 and Supplementary Data. Public genome assemblies and RNA-seq datasets were retrieved from AlliumDB, NABIC, NCBI-SRA and the Onion Genome Database under the accession numbers and BioProjects indicated in the corresponding supplementary tables.

## Abbreviations

A: Net photosynthetic rate aa, amino acids
AQP(s): aquaporin(s)
ACE: apparent carboxylation efficiency
Ci: internal CO_2_ concentration
E: transpiration rate
ENA: European Nucleotide Archive FW, fresh weight
Gs: stomatal conductance
MIP(s): membrane intrinsic protein(s)
NCBI: National Centre for Biotechnology Information
NIP(s): nodulin 26-like intrinsic protein(s)
PAR: photosynthetically active radiation PCA, Principal Component Analysis
pI: isoelectric point
PIP(s): plasma membrane intrinsic protein(s) RIN, RNA Integrity Number
RNA-seq: RNA sequencing ROS, reactive oxygen species
SIP(s): small and basic intrinsic protein(s)
SRA: Sequence Read Archive database
TIP(s): tonoplast intrinsic protein(s)
VST: variance stabilizing transformation WUE, water use efficiency
XIP(s): X-intrinsic proteins(s)

## Notes

### Competing Interest Statement

The authors have declared no competing interest.

